# Quantifying Heterogeneity in the Genetic Architecture of Complex Traits Between Ethnically Diverse Groups using Random Effect Interaction Models

**DOI:** 10.1101/421149

**Authors:** Yogasudha Veturi, Gustavo de los Campos, Nengjun Yi, Wen Huang, Ana I. Vazquez, Brigitte Kühnel

## Abstract

In humans, most genome-wide association studies have been conducted using data from Caucasians and many of the reported findings have not replicated in other populations. This lack of replication may be due to statistical issues (small sample size, confounding) or perhaps more fundamentally to differences in the genetic architecture of traits between ethnically diverse subpopulations. What aspects of the genetic architecture of traits vary between subpopulations and how can this be quantified? We consider studying effect heterogeneity using random-effect Bayesian interaction models. The proposed methodology can be applied using shrinkage and variable selection methods and produces useful information about effect heterogeneity in the form of whole-genome summaries (e.g., SNP-heritability and the average correlation of effects) as well as SNP-specific attributes. Using simulations, we show that the proposed methodology yields (nearly) unbiased estimates of genomic heritability and of the average correlation of effects between groups when the sample size is not too small relative to the number of SNPs used. Subsequently, we used the proposed methodology for the analyses of four complex human traits (standing height, high-density lipoprotein, low-density lipoprotein, and serum urate levels) in European-Americans (EAs) and African-Americans (AAs). The estimated correlations of effects between the two subpopulations was well below unity for all the traits, ranging from 0.73 to 0.50. The extent of effect heterogeneity varied between traits and SNP-sets. Height showed less differences in SNP effects between AAs and EAs whereas HDL, a trait highly influenced by life-style, exhibited greater extent of effect heterogeneity. For all the traits we observed substantial variability in effect heterogeneity across SNPs, suggesting it varies between regions of the genome.

---

**P**opulation structure is a pervasive feature in plant, animal, and human populations (Gaggiotti *et al.* 2009; Pfenninger *et al.* 2011; Puckett *et al.* 2014). In population genetics, differentiation between subpopulations is often measured by comparing allele frequencies, e.g., using the ‘F-statistic’ (Malécot 1947; Wright 1949) and (Cockerham 1969). In genome-wide association studies (GWAS), population differentiation is predominantly viewed as a confounder (Astle and Balding 2009) that can lead to spurious associations (Lander and Schork 1994; Deng 2001; Marchini *et al.* 2004; Liu *et al.* 2011). To address this problem a variety of methods have been proposed (Price *et al.* 2010). However, rather than a confounder, population stratification can act as an effect-modifier, leading to heterogeneity in the genetic architecture of traits.

The evolutionary dynamics involved in the processes that lead to population structure can result in subpopulations with heterogeneity in allele frequencies and linkage disequilibrium (LD) patterns (Gabriel 2002). Moreover, in some instances, ethnic background correlates with environmental exposures (e.g., diet, income, lifestyle) and this can lead to genetic-by-environment interactions. All these differences between ethnic groups can induce heterogeneity in the genetic architecture of traits (de los Campos and Sorensen 2014). Quantifying the extent of effect heterogeneity between ethnically diverse groups is relevant across disciplines and can shed light on whether results obtained in one group are expected to replicate in others. This is particularly important when we consider that the vast majority of GWAS have been conducted using data from Caucasians and that results reported from these studies do not always replicate in other populations, which may indicate differences in genetic architectures between ethnic groups (Greene *et al.* 2009a; Kraft *et al.* 2009; Ng *et al.* 2014).

Several studies have demonstrated (or alluded to) effect heterogeneity between ethnic groups (Ntzani *et al.* 2012; de Candia *et al.* 2013; Li and Keating 2014; Brown *et al.* 2016). Most of these studies measured effect heterogeneity by estimating the average correlation of marker effects between two or more ethnically diverse groups.

One may attempt to estimate effect correlations by quantifying the average correlation of estimated effects from GWAS conducted in different ethnic groups. However, estimation errors make the simple correlation of estimates of effects a seriously biased (towards zero) estimate of the correlation of (true) effects (see Appendix C for a simple demonstration of this). To overcome this problem, several studies have used multivariate Gaussian random regression models. Such methods have been considered in both animal and plant breeding (Wei and Werf 1994; García-Cortés and Toro 2006; Karoui *et al.* 2012; Olson *et al.* 2012; Christensen *et al.* 2014; Lehermeier *et al.* 2015) as well as in human genetics (e.g. de Candia et al., 2013; Lee et al., 2012). Another approach estimates the correlation of effects using an extension of the LD-score regression (Brown *et al.* 2016).

The methods above described provide whole-genome summaries such as SNP-heritability and average correlation of effects. However, they don’t shed light on how effect heterogeneity may vary across regions of the genome or between SNP sets. Moreover, the random regression methods commonly used to estimate the average correlation of effects make the Gaussian assumption; therefore, they cannot be used with priors that induce differential shrinkage of estimates (e.g., double-exponential (Park and Casella 2008) or a combination of shrinkage and variable selection (e.g. spike slab, Ishwaran and Rao 2005)). To overcome this limitation, we consider modeling effect heterogeneity using a Bayesian random-effect interaction model that decomposes SNP effects into main and interaction components. Unlike previously used methods, the proposed approach can be applied with both shrinkage and variable selection priors and offers both whole-genome and SNP-specific measures of effect heterogeneity. In addition to estimating average correlation of effects, the proposed method can also estimate the proportion of non-zero effects in the considered SNP set; we will consider both measures when quantifying effect heterogeneity.

Using simulations, we demonstrate that the proposed method yields nearly unbiased estimates when sample size (n) is not too small relative to the number of markers (P) used. Subsequently, using data from the multi-ethnic Atherosclerosis Risk in Communities (ARIC) study, we applied the proposed method to conduct joint analyses of individuals of European and African ancestry [hereinafter referred to as European-Americans (EAs) and African-Americans (AAs), respectively] for traits with varying genetic architectures. These subpopulations have important differences in allele frequencies, LD decay (Shifman 2003) and cultural and socio-economic factors that are linked to environmental exposures.

Our results show that for the four traits there is a large extent of effect heterogeneity with average correlations of effects well below one. However, the extent of effect heterogeneity between EAs and AAs also varies between traits (the correlation of effects was highest for height and lower for lipid traits). Moreover, we show that for HDL, LDL and serum urate there is great deal of variability in effect heterogeneity between SNPs with many SNPs showing almost no effect heterogeneity and others exhibiting substantial effect differences between AAs and EAs.

## Materials and Methods

Meuwissen et al. (2001) proposed to analyze and predict complex traits by regressing phenotypes on whole-genome panels of SNPs. Their model was developed with reference to a homogeneous population. Here, following (de los Campos *et al.* 2015), we consider extending the whole genome regression model by including random-effect interactions between markers and groups.

Considering two groups, the regression of phenotypes 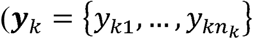, where *k*= 1,2 indexes groups and *n*_*k*_ denotes the number of individuals in the *k*^*th*^ group) on *p* markers (e.g., SNPs) can be represented as follows:
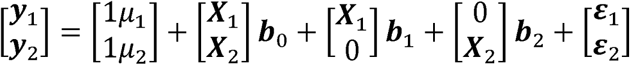
(1)where *µ*_1_ and *µ*_2_ are group-specific intercepts, 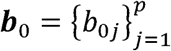 is a vector of ‘main effects’, 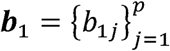 and 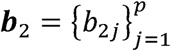 are group-specific interactions and 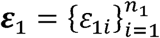 and 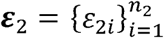 are error terms. In our models, we assume IID Gaussian errors with group-specific variances that is 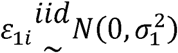 and 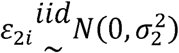.

**Marker effects** in groups 1 and 2 are defined by the sum of the main and group-specific terms, that is, *β*_1*j*_ = *b*_0*j*_ + *b*_1*j*_ and *β*_2*j*_ = *b*_0*j*_ + *b*_2*j*_, respectively. Since the number of markers is usually large relative to sample size we treat both main and interaction effects as random. Depending on the distribution assigned to SNP effects the model can induce variable selection, shrinkage, or a combination of both (Ishwaran and Rao 2005; Gianola *et al.* 2009; de los Campos *et al.* 2013). To illustrate, we considered two priors for main and interaction effects: a Gaussian distribution and a prior with a point of mass at zero and a Gaussian slab, also known as BayesC (Habier *et al.* 2011).

In the **Gaussian setting** we assign independent Normal priors with null mean and with different variances for the main and interaction effects, that is

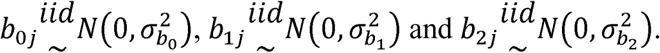

Above, 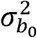, 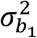 and 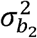 represent the prior variances of the main and interaction effects, respectively.

For the **Spike-Slab prior** we adopt the assumptions of model BayesC (Habier *et al.* 2011), with set-specific variances and proportions of non-zero effects, that is
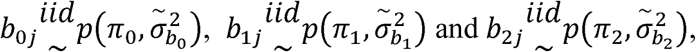
where 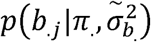 is a mixture distribution of the form 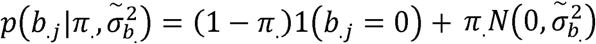, for 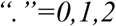. Here, *π* represents the proportion of non-null effects.

**Hyper-parameters**. In the Gaussian model the hyper-parameters are the error variance and the three variances of effects, that is 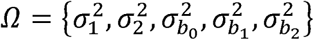. In BayesC the hyper-parameters also include the proportion of non-null effects; therefore: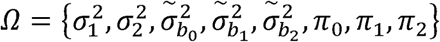.

These parameters control the extent of shrinkage and variable selection and how the architecture of effects may vary between groups. In our implementation, we treat them as unknown and therefore assign prior distributions to them. For variance parameters, the conjugate prior is the scaled-inverse chi-squared. However, this prior can have some influence on inference. Therefore, instead we use a prior for variance parameters that is a transformation of the beta distribution (Appendix A). For the proportion of non-zero effects {π_0_,π_1_,π_2_} we use independent identical beta priors. This allows us to accommodate different effect distributions for different traits and sets of SNPs. Further details about this are provided in the Data Analysis section below.

The models described above can be used to estimate several parameters that are descriptive of the trait architecture. Whole-genome summaries of the trait architecture and of effect heterogeneity include *the proportion of variance explained by SNPs* (or genomic heritability, e.g., de los Campos et al., 2014) in each of the ethnic groups, the *average correlation of effects* and the *average proportions of non-zero effects* (either main effects, interaction terms or total effects). Samples from the posterior distribution can also be used to estimate SNP-specific parameters such as the posterior correlation of a SNP effect, *ρ*_*j*_ = *Cor*(*β*_1*j*_, *β*_2*j*_).

Genomic variance, genomic heritability and the average correlation of effects were estimated using the methods described by Lehermeier et al. (2017). Briefly, at each iteration of an MCMC algorithm, we used the samples of the main and interaction effects to form marker effects, *β*_1*j*(*s*)_ = *b*_0*j*(*s*)_ + *b*_1*j*(*s*)_ and *β*_2*j*(*s*)_ = *b*_0*j*(*s*)_ + *b*_2*j*(*s*)_ (here, *s=1,…,N* is an index for the N samples collected). A sample from the posterior distribution of the correlation of effects was obtained from *ρ*_*s*_ = *Cor*(*β*_1*j*(*s*)_,*β*_2*j*(*s*)_) where *Cor*( ) represents Pearson’s product moment correlation.

Likewise, at each iteration genomic values can be obtained from ***u***_1(*s*)_ = ***X***_1_*β*_1(*s*)_ and ***u***_2(*s*)_ = ***X***_1_*β*_2(*s*)_. Therefore, a sample for the posterior distribution of the genomic variances for each group was computed as 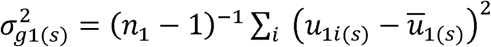 and 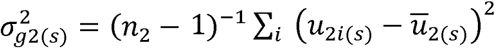, where 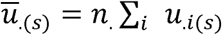. Finally, samples from the posterior distribution of genomic heritability were obtained using the following: 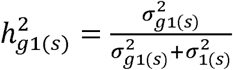 and 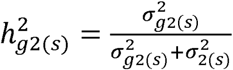.

### Data

Our simulation and real data analyses were based on data from the Atherosclerosis Risk in Communities (ARIC) study. ARIC is a prospective epidemiologic study sponsored by the National Heart, Lung, and Blood Institute (NHLBI) conducted in four U.S. communities to study the causes of atherosclerosis and other Cordiovascular risk factors such as blood lipids, lipoprotein cholesterols, and apolipoproteins. It has a total sample size of 15,792 AA and EA men and women aged 45-64. A total of 13,113 individuals were genotyped using the Affymetrix array with a total of 934,940 SNPs. Genotype and phenotype data from the ARIC study was acquired through the dbGaP (IRB number 15-745; r050661 and study accession number phs000280.v1.p1).

### Genotypes

We retained SNPs that had minor allele frequency higher than 1% in at least one of the two ethnic groups (SNPs that are monomorphic in only one of the two groups are clearly heterogeneous), had higher than 95% calling rate, and were mapped to one of the 23 human chromosomes. After QC, we retained 828,822 SNPs. Individuals with a 5% missing rate in their genotypes were removed. Individuals were classified as EA or AA based on self-reported ethnicity. We confirmed this assignment by deriving principal components from the genomic relationship matrix (Figure S1). Subsequently we computed genomic relationships within each ethnic group and removed those that had within-group genomic relationships higher than 0.075; this ensured that we only retained distantly related individuals. After standardization to an average diagonal value of 1, the diagonal elements of the marker-derived genomic relationship matrix ranged between 0.85 and 1.15. Thus, the final data sets comprised only distantly related individuals including 6,627 EAs and 1,601 AAs.

### Simulations

We simulated phenotypes using genotype data from the ARIC study from 6,627 EAs and 1,601 AAs. Phenotypes were simulated under an additive genetic model with a heritability of 0.5 for both groups. We considered scenarios with the number of markers (n) varying from 100 to 10,000 and the true correlations of effects between groups varying from 0.2 to 0.8. In a first simulation setting we assumed that all the markers had effects on both groups. In a second setting, we assumed that 50% of the loci had effects on both groups, 20% had effects on EAs but not on AAs, 20% had effects on AAs but not on EAs and 10% had no effects on either group (non-causal variants). Further details of the simulation are given in Appendix B.

### Analyses of Four Complex Human Traits

For our real-data analyses we considered four complex phenotypes: human height (cm), HDL (mmol/L) and LDL (mmol/L) cholesterol and serum urate (mg/dL). Individuals with height < 147 cm, LDL > 10 mmol/L and serum urate > 15 mg/dL were removed. We did not identify clear outliers for HDL. Transformation of the traits was not considered necessary (Figure S2). Phenotypes were pre-corrected for ethnicity, age and sex.

Models were fitted to subsets of SNPs selected based on single-marker regression (GWAS) p-values derived from data that did not include ARIC. For height, GWAS p-values were derived from the full release of the UK Biobank. For HDL and LDL p-values were from the Global Lipids Genetics Consortium (GLGC) computed after excluding data from ARIC. Finally, for serumurate, p-values were from the Global Urate Genetics Consortium (GUGC), also derived without using data from ARIC. The simple ranking of markers based on association *p*-values would lead to sets of highly redundant markers, i.e., markers in high LD (see Figure S3). To avoid this, we designed a windows-based selection algorithm where a window was defined as a set of consecutive SNPs that exceeded a given –log_10_(*p*-value) cutoff (this was done on a per-trait basis). The goal of this method was to avoid oversampling SNPs from genomic regions of high LD. Windows were made on a per-trait basis at –log_10_(*p*-value) cutoffs of 2, 2.3, 2.6, 3, 5, and 8 (Table S1). SNPs that cleared a given –log_10_(*p*-value) cutoff were termed “significant” at that cutoff (See Figure S4).

We fitted the interaction model to each of the four traits and each of the SNP-sets above described. As part of sensitivity analyses, we also fitted the same models to randomly chosen sets of SNPs (of sizes 500, 1,000, 2,500, 5,000, and 10,000 SNPs, respectively). Finally, we also evaluated the estimates obtained within the EA ethnic group label randomly permuted.

### Software

Models were fitted using a modified version of the BGLR (Pérez and de los Campos 2014) R package (available at: https://github.com/gdlc/BGLR-R and at https://cran.r-project.org/web/packages/BGLR/index.html) that implements a weakly informative prior for variance parameters based on a transformation of the beta distribution (de los Campos *et al.* 2009) described above. We ran the Monte Corlo Markov Chain algorithm for 45,000 iterations; the first 15,000 iterations were discarded as burn-in and the remaining samples were thinned at a thinning interval of 5.

**Hyper-parameters**. BGLR assigns a beta prior to the proportion of non-zero effects, we choose the shape parameters of the beta prior to be equal to 1, which gives a uniform prior in the 0-1 interval. For variance parameters we devised a prior which is a modified version of the Beta prior (see Appendix A) and used shape parameters equal to 1.01 to obtain an almost uniform prior for variance parameters within the interval [0,K] where K was twice the variance of the phenotype.

### Data availability

Supplemental files are available as a zipped folder (not on FigShare). File S1 contains supplementary figures, tables and Appendices. The IRB number for ARIC dataset is 15-745; r050661 and the study accession number for ARIC dataset: phs000280.v1.p1.

## Results

### Simulations

In both simulation settings, the <u>*genomic heritability*</u> was estimated with almost no bias using both Gaussian and BayesC priors (see Figures 1 and S5 for the first and second simulation scenarios, respectively). The standard errors were higher for AAs as compared to EAs, which was expected given that the sample size was smaller for AAs. As one would expect, the standard errors also increased with the number-of-loci/sample-size ratio. Using the BayesC prior, the estimates of genomic heritability were mildly biased across all values of true effect correlation when the number of QTL was greater than 10,000. There was a mild downward bias when the true correlation was very low (0.2 or 0.4) and there was a mild upward bias when the true correlation was high (0.8).

**Figure 1.**
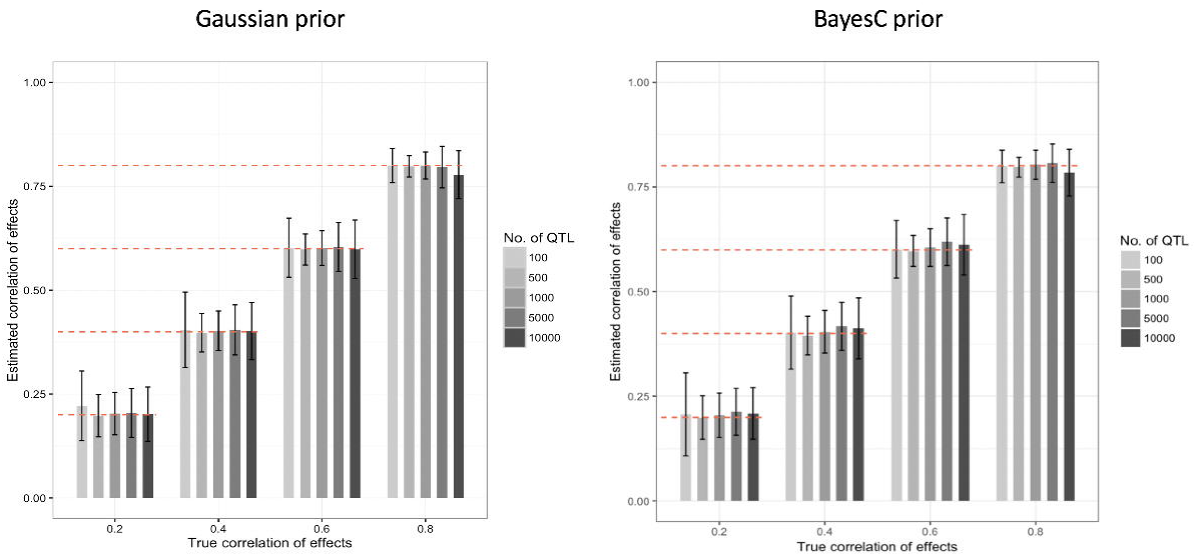
Average estimates of genomic heritability obtained in the first simulation scenario, by prior and number of SNPs used. The simulated heritability was 0.5, bars represent the average estimates over 200 Monte Corlo replicates and the vertical lines gives +/- standard errors. Results for the 2^nd^ simulation scenario are presented in Figure S5.

Estimates of <u>*effect correlations*</u> were also nearly unbiased (see Figures 2 and S6). However, the standard errors were very large, particularly when the correlations were low. In scenarios involving more than 5,000 QTL we observed a small downward-bias. Likewise, we observed a small upward bias for the Gaussian prior when the number of QTL was 100 (Figure 2 and Figure S6). The average standard error of the estimated correlation was high with the smallest (100) and the largest (10,000) numbers of QTL and lower for scenarios in between.

**Figure 2.**
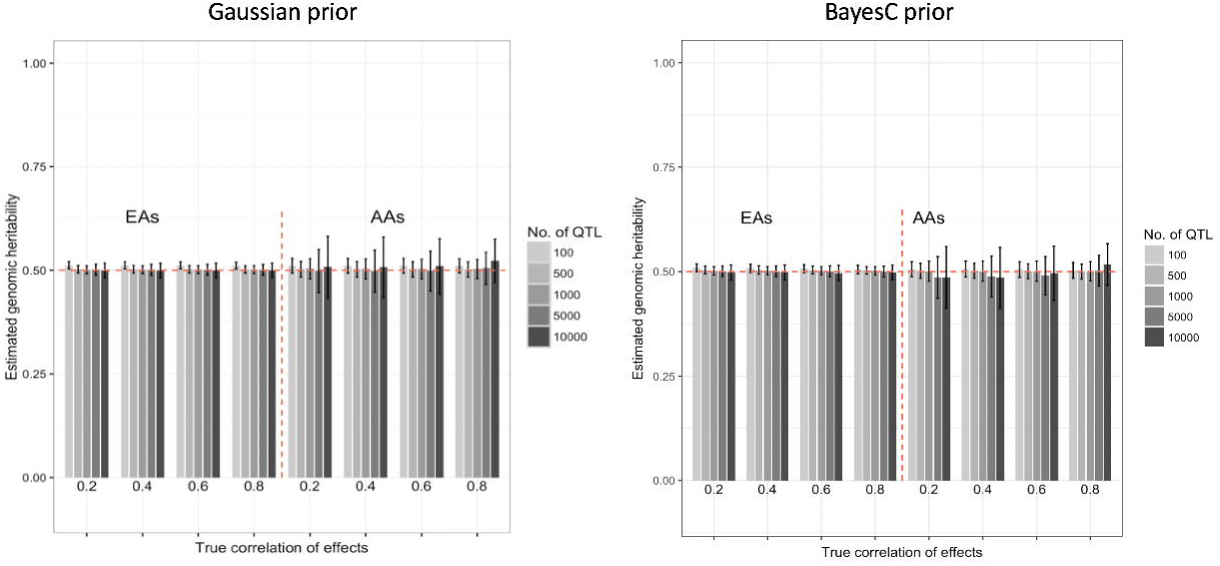
Average estimates of the correlation of effects in the first simulation scenario by prior and number of SNPs used. The simulated heritability was 0.5; bars represent the average estimates over 200 Monte Corlo replicates and the vertical lines gives +/- standard errors. Results for the 2^nd^ simulation scenario are presented in Figure S6.

### Analyses of Four Complex Human Traits

Since our simulations revealed that n/P ratio of at least 1/3 results in nearly unbiased estimates of genomic heritability, we fit our model to subsets of markers instead of using whole-genome data (see methods for a description of how these subsets were obtained). Figure 3 shows the estimated <u>*genomic heritability*</u> obtained using the BayesC prior, by trait, ethnicity and the set of SNPs used. (The results obtained with the Gaussian prior are displayed in Figure S7). As expected, the estimated genomic heritability increased with the number of SNPs used. Interestingly, this parameter was systematically higher in EAs than in AAs for height and HDL, and the order was reversed in other traits (LDL and serum urate). However, the credibility intervals between both ethnic groups overlapped for all traits except height. The genomic heritability estimates obtained with the Gaussian prior were similar to the ones found with the BayesC prior (see Figure S7) for all traits except serum urate, which yielded larger estimates for AAs than those obtained using the BayesC prior.

**Figure 3.**
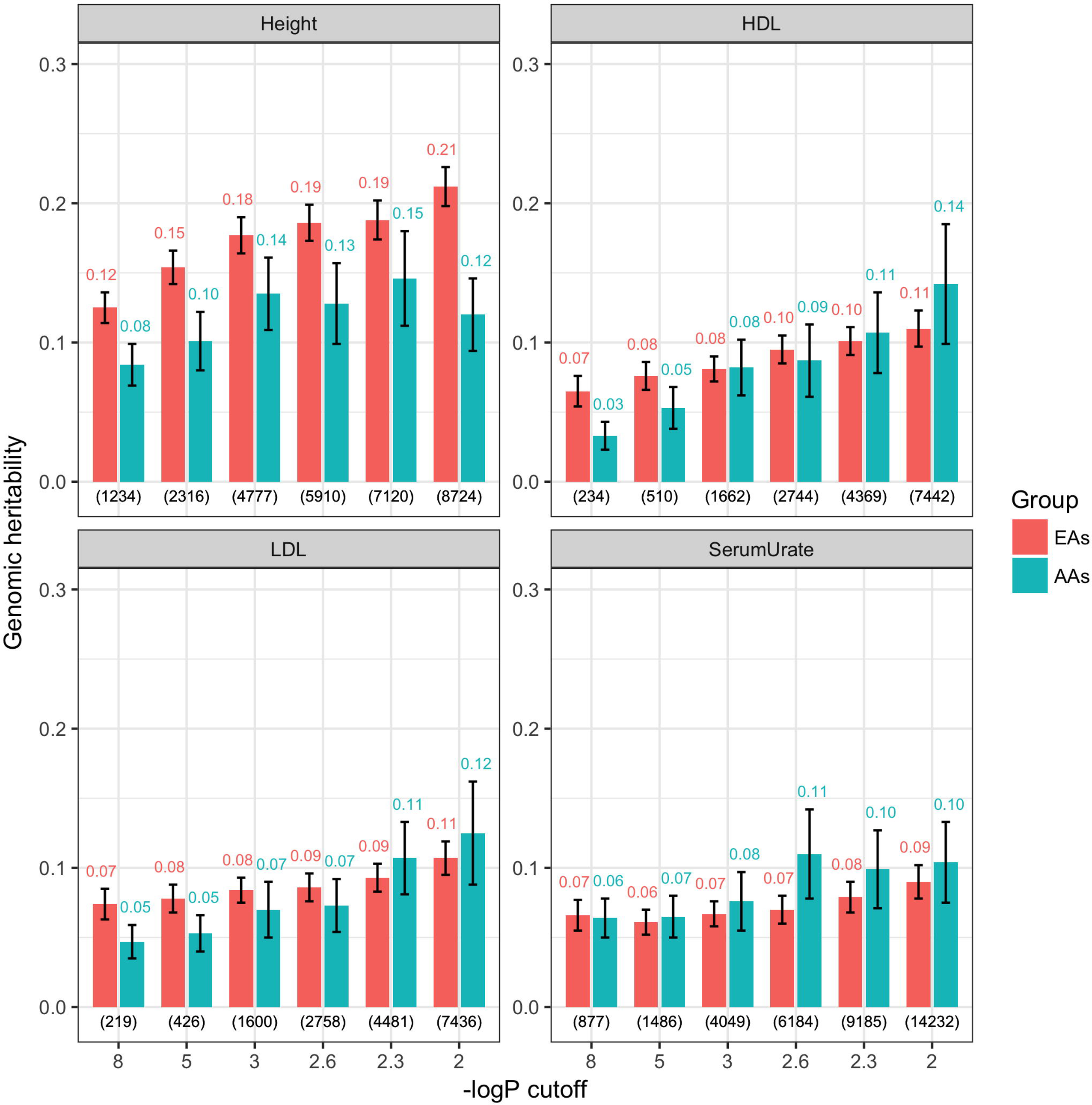
Proportion of variance explained by subsets of SNPs obtained with the BayesC-interaction model, by trait, ethnicity and SNP set. Estimated (median) genomic heritability (y-axis) is plotted by trait, ethnicity and log_10_(*p*-value) cutoff used to choose markers from GWAS consortia (excluding ARIC). Numerals above the bars indicate the proportion of variance explained by either ethnic group and the corresponding number of SNPs used for model fitting (in parentheses at the bottom). Vertical lines give estimates of +/- posterior standard deviation.

Figure 4 shows the estimated <u>*correlation of effects*</u> between AAs and EAs obtained with the BayesC prior, by the set of SNPs used in the analysis. The estimated average correlation of effects ranged from 0.711 (for height with the SNP-set obtained with –log_10_(*p*-value) cutoff of 8) to 0.500 (for HDL with the SNP-set obtained with a –log_10_(*p*-value) cutoff of 2.3). Overall the correlation of effects was highest for height and serum urate and lowest for LDL and HDL. In all traits except HDL the correlation of effects tended to decrease as more SNPs were added in the model; however, the confidence regions for the different SNP sets overlapped. The estimated correlation of effects with the Gaussian prior for marker effects was similar to those obtained using the BayesC prior, with subtle differences between the two priors for height, HDL and LDL (Figure S8).

**Figure 4.**
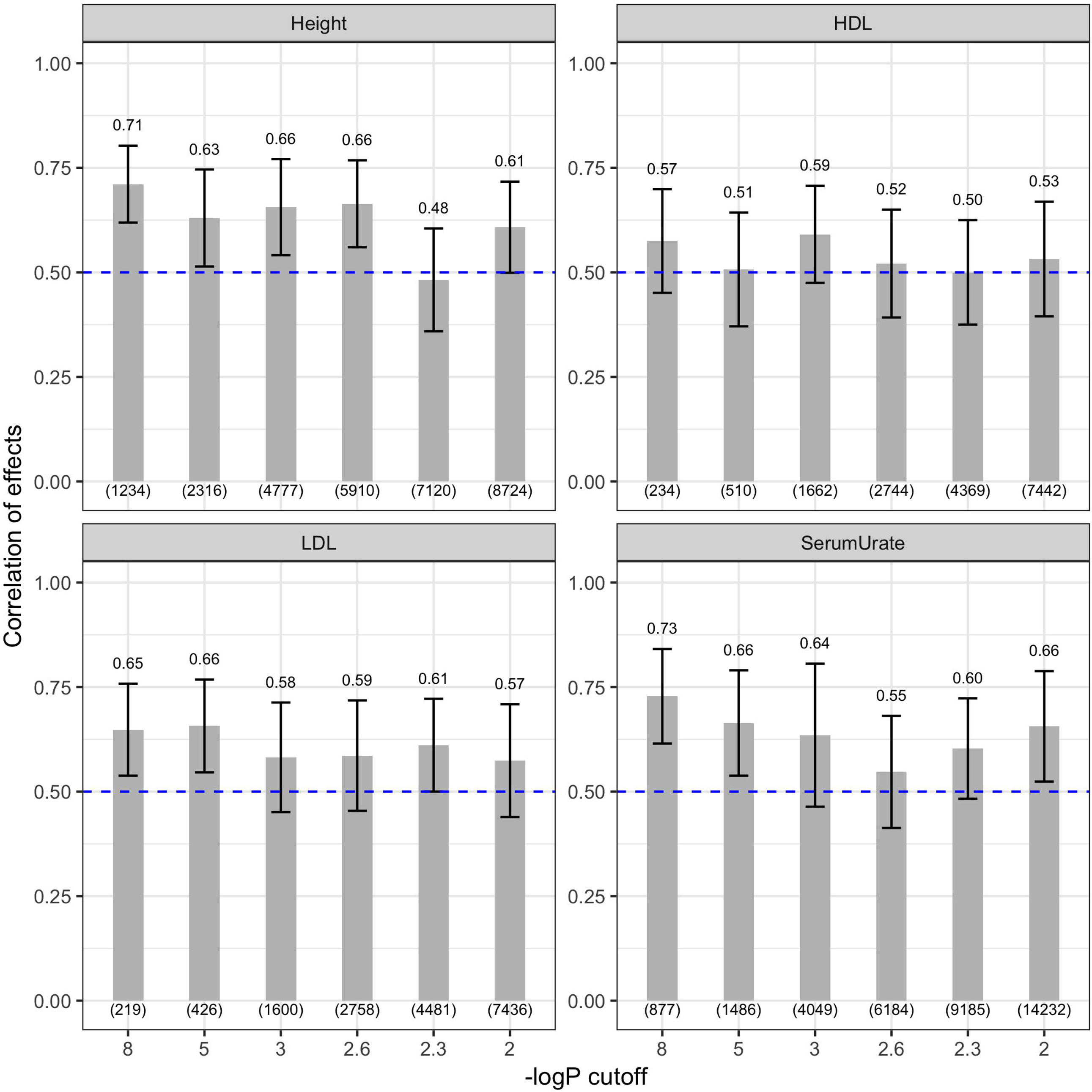
Estimated correlation of effects between African Americans (AAs) and European Americans (EAs) obtained with the BayesC-interaction model, by trait and SNP set. Estimated correlation of effects between AAs and EAs (y-axis) is plotted by trait using markers selected from GWAS consortia (excluding ARIC). In each plot, the numerals above the bars indicate the median correlation of effects and the number of SNPs used for model fitting (in parentheses at the bottom). Vertical lines give estimates of +/- posterior standard deviation.

Figure 5 shows the estimated proportion of non-zero SNP effects obtained with the BayesC prior, by trait, ethnic group and SNP-set. For both groups, the proportions of non-zero effects were high at large –log_10_(*p*-value) cutoffs and decreased as the number of markers included in the model increased. For height, the proportions of non-zero effects were similar between EAs and AAs. However, for LDL (and serum urate to a lesser extent) the decrease in the proportion of non-zero effects was stronger in EAs. Figure S9 displays the proportion of non-zero main and interaction effects. The proportion of non-zero main effects decreased as the number of SNPs increased and the proportion of non-zero interaction terms tended to remain constant (except for the LDL-interactions for EAs). Interestingly, the proportion of non-zero effects dropped very fast with the number of SNPs for HDL, LDL and serum urate, suggesting for these traits that the effect architecture may be highly heterogeneous between AAs and EAs.

**Figure 5.**
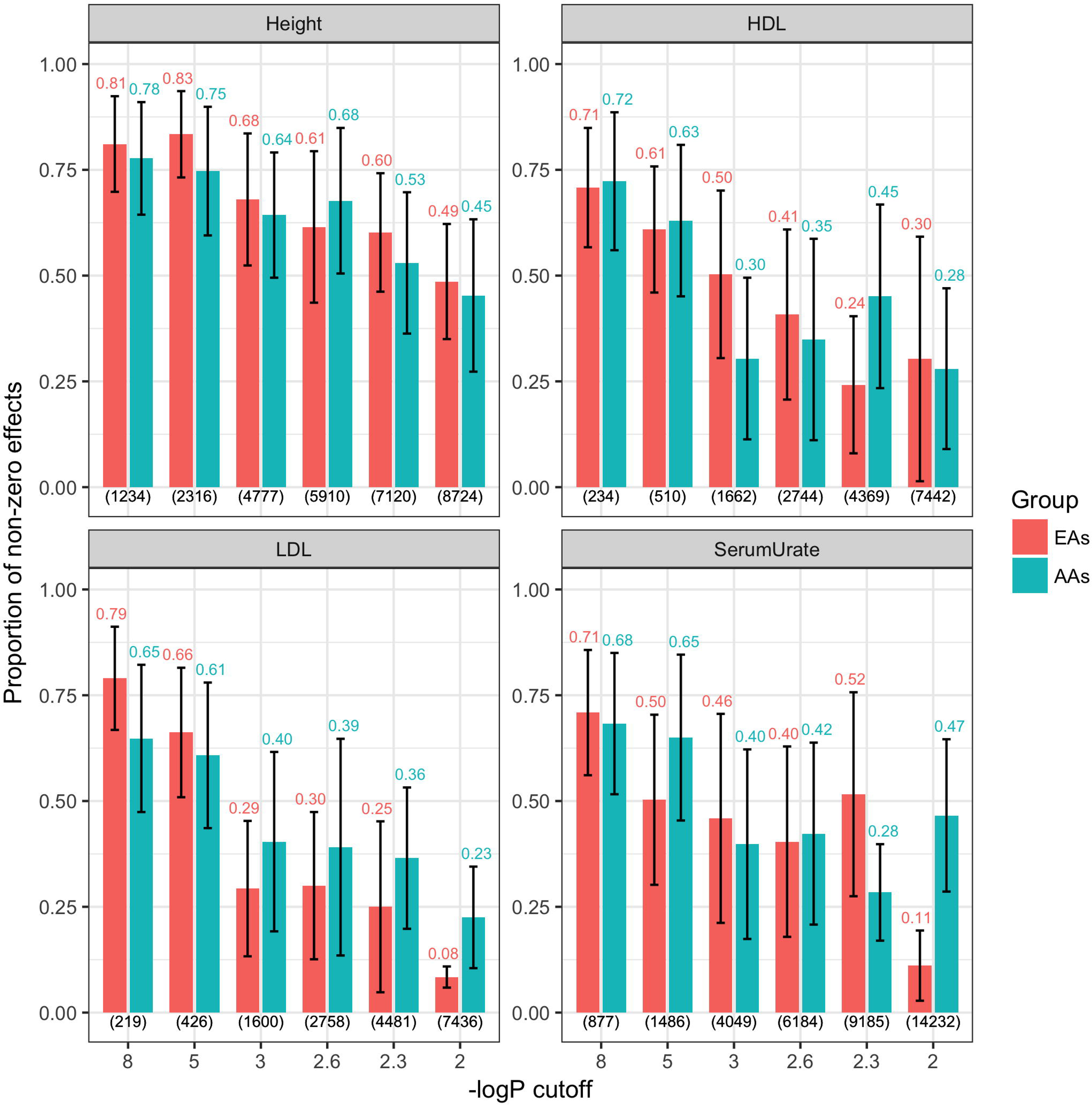
Estimated proportion of non-zero effects between African Americans (AAs) and European Americans (EAs) obtained with the BayesC-interaction model, by trait and SNP set. Estimated proportion of non-zero effects between AAs and EAs (y-axis) is plotted by trait using markers selected from GWAS consortia (excluding ARIC) at 6 different –log_10_(*p*-value) cutoffs. In each plot, the numerals above the bars indicate the proportion of non-zero effects obtained using either ethnic group and the corresponding number of SNPs used for model fitting (in parentheses at the bottom). Vertical lines give estimates of +/- posterior standard deviation.

Figures 3-5 (and the corresponding supplementary figure S9), correspond to whole-genome summaries (SNP-heritability, average correlation of effects, proportion of non-zero effects). However, the models also render SNP-specific summaries. Figure 6 shows the posterior mean of the correlation of effects between ethnic groups for individual SNPs by trait for the SNP set obtained using a –log_10_(*p*-value) cutoff of 2. We had no SNP with negative posterior correlation of effect. For height, the posterior correlation of individual-SNP effects ranged from 0.4-0.8. However, for HDL, LDL and serum urate, there was more variability among SNPs, with several SNPs having posterior correlation of effects greater than 0.8 and many with posterior correlation of effects smaller than 0.4.

**Figure 6.**
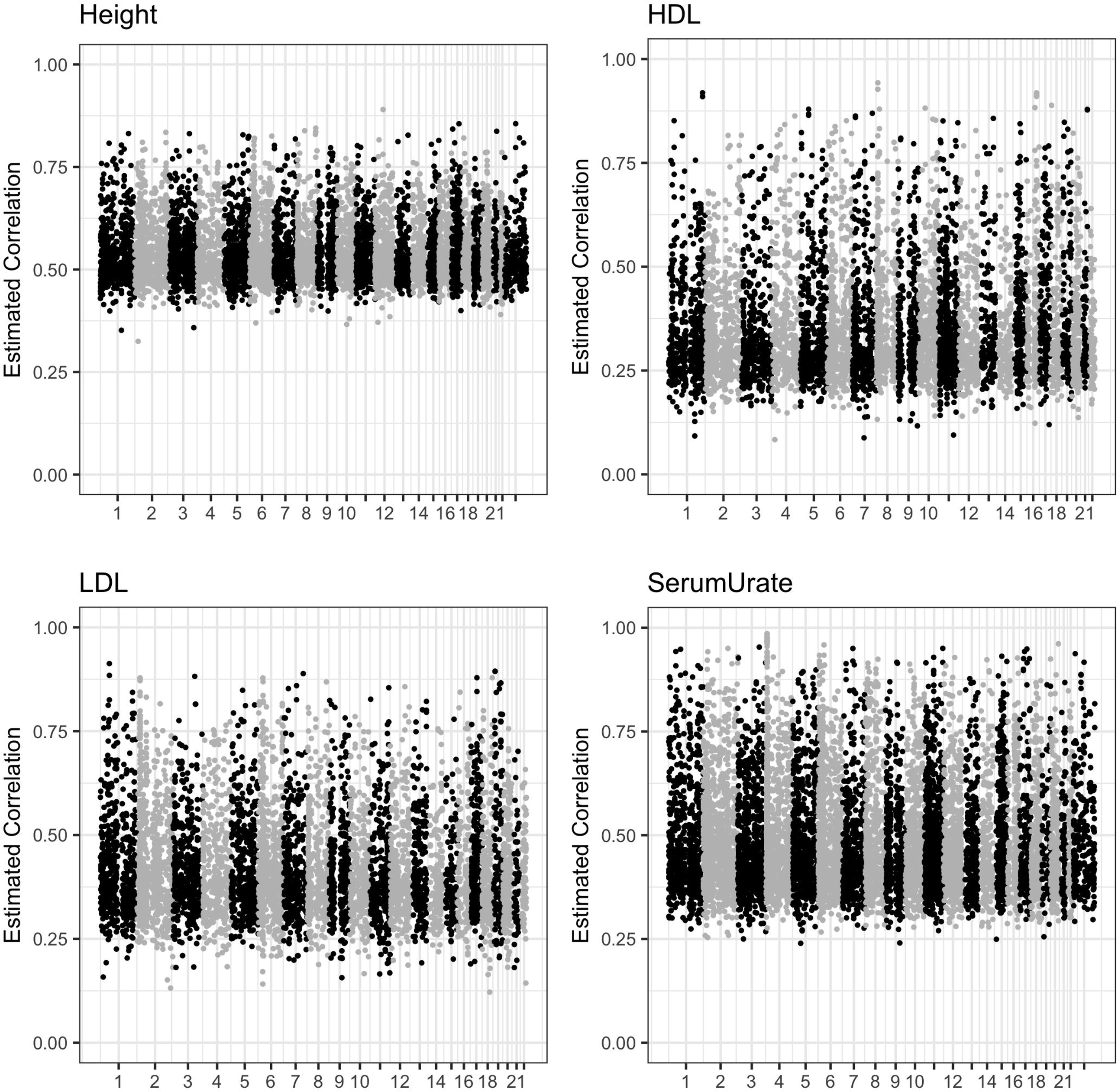
Posterior correlation of individual SNP effect between African Americans (AAs) and European Americans (EAs), by trait for SNPs that clear a –log_10_(*p*-value) of 2. Plots are categorized by trait and in each plot, the estimated effect correlation of individual SNP effects (y-axis) is plotted against chromosome number (x-axis).

## Discussion

Genome-Wide Association (GWA) studies have discovered large numbers of variants associated with important traits and diseases in humans (Welter *et al.* 2014) but have been conducted predominantly in Caucasian populations (Haga 2010; Rosenberg *et al.* 2010). Although more recent works have recognized inclusion of diverse ethnic groups, especially African Americans (e.g. Brant et al., 2017; Park et al., 2017; Taylor et al., 2016) the total number of GWAS studies for African Americans is still fairly low compared to populations of European ancestry (Peprah *et al.* 2015) and replication of signals in African American populations is much less common (Marigorta and Navarro 2013). Moreover, the associations reported to be strong in Caucasians have been weaker (or even non-significant) in other ethnic groups (Gudbjartsson *et al.* 2007; Omori *et al.* 2008; Yamada *et al.* 2009; Barnholtz-Sloan *et al.* 2011; Tsai *et al.* 2014; Prasad *et al.* 2017) and some studies have reported effects with opposite sign in different populations (Lewis *et al.* 2008; Yamada *et al.* 2009). Other recent studies have also indicated presence of genetic heterogeneity between ethnic groups for different quantitative/binary traits (Brown *et al.* 2016; de Vlaming *et al.* 2017; Zhou *et al.* 2018). While some of these differences could be attributed to dissimilarities in LD and sample sizes (e.g. some well-powered studies have shown strong overlaps in GWAS-significant variants between Europeans and other ethnic groups (Franceschini *et al.* 2013; Okada *et al.* 2014)), there could also be novel variants specific to the ethnic group (Chen *et al.* 2017), indicating that genetic heterogeneity could depend on other genetic factors and be trait-specific. Understanding the reasons that underlie these differences and quantifying the degree of similarity in the architecture of a trait across populations represents an important research goal.

A naïve approach for assessing the degree of similarity in the genetic architecture of a trait between two groups would be to simply correlate estimated effects obtained from stratified analyses (e.g., GWAS conducted within each group). However, errors in estimates of effects make this estimator highly biased towards zero (see Appendix C). Our empirical results verified this (Table S2): the sample correlation of estimated effects was consistently lower than the one estimated using variance components (a nearly unbiased estimator from our simulations). Our results also showed that the difference between the naïve estimator of the correlation and the one based on variance components increases as more small-variance markers are used in the model.

In humans, Brown et al. 2016 considered quantifying the average correlation of effects using summary-based association statistics. Their approach extended LD score regression (Bulik-Sullivan *et al.* 2015) to multiple ethnic groups and has the advantage, relative to the method discussed here, that it can be used with summary statistics. However, some authors have questioned the assumptions of the LD score regression method (Speed *et al.* 2018) and accurate estimation requires using several thousands of SNPs. Thus, the method is not well-suited for studying effect similarity within genomic regions, something that the proposed method can achieve. Also, the LD-score method requires access to good quality external reference panels (especially for non-European populations) to construct population LD matrices, assumes normal distribution of marker effects (infinitesimal model) and is not directly applicable to admixed populations (or populations that have long range LD).

In this study, we propose to study ethnic differences in the architecture of traits using a random effect Bayesian interaction model. The proposed approach can be used to estimate whole-genome summaries such as (a) the proportion of variance explained by SNPs, (b) the average effect correlation, (c) proportion of non-zero effects, as well as finer features of the trait architecture (e.g. SNP-specific correlation of effects). Similar approaches have been considered in animal and plant breeding (e.g., Christensen et al., 2014; García-Cortés and Toro, 2006; Lehermeier et al., 2015; Wei and Werf, 1994) and in human genetics (e.g., de Candia et al., 2013; Lee et al., 2012) for the analysis of data from heterogeneous populations. However, previous studies were based on Gaussian assumptions and only offered whole-genome summaries of the trait architecture. The approach presented in this study is more flexible in that it can be used with both shrinkage and variable selection priors (Ishwaran and Rao 2005; Park and Casella 2008) and can be used to infer not only whole-genome features but also regional and SNP-specific features of the trait architecture.

Among many parameters, the proposed method yields an estimate of effect correlation. This parameter, should not be confounded with the classical concept of genetic correlation (Reeve 1953). While the former provides a measure of the extent of effect similarity/heterogeneity between two groups for the same trait, the latter is defined between two traits for the same individual. Moreover, the forces that contribute to the genetic correlation between traits (pleiotropy and LD between alleles at causal loci affecting different traits) are different from the forces that may contribute to effect heterogeneity between ethnic groups (e.g., differences in allele frequency and genetic by-environmental interactions). Thus, genetic correlation and correlation of effects between ethnically diverse groups should be treated as conceptually different.

We evaluated the proposed methodology under two different priors (Gaussian and BayesC) using simulations and applied it to real human data to study the genetic architecture of four traits (Height, HDL, LDL, Serum urate) in EAs and AAs. Our simulation study revealed that both Gaussian and BayesC priors yield nearly unbiased estimates of genomic heritability and effect correlations. From our real data analyses, we observed similar proportions of variance explained with both BayesC and Gaussian priors for marker effects (Figure 3 and S7). Except height, the average proportions of variance explained across all marker sets were similar between EAs and AAs. For height, the average proportion of variance explained was greater among EAs than among AAs. This is likely due to the fact that the SNPs used for the analysis of height were selected using GWAS results entirely based on data from Caucasians (UK Biobank); the same trend was not observed for other traits perhaps because there was some mixture in ethnicity in the other GWAS consortia from which markers were chosen (GLGC, GUGC). When we fit similar models using randomly chosen markers (Figure S10), we observed that the proportion of variance explained by randomly selected markers was smaller than that explained by regression on markers selected from GWAS results for both EAs and AAs. This showed that indeed, selection based on GWAS results leads to more informative markers in both populations.

Our analyses also revealed important differences in correlation of effects between traits. The **estimated correlation of effects** ranged from 0.482-0.728, indicating the presence of genetic heterogeneity across all four traits, even for strongly associated markers (Figure 4). For height the correlation of effects was highest when using SNPs that had the smallest GWAS p-value (likely SNPs with relatively large effect and not very extreme allele frequency), suggesting that the correlation of effects may be lower for SNPs with small effects and those with extreme allele frequencies.

Height had higher correlation of effects than serum urate and lipid traits, suggesting that height may have a more similar genetic architecture between EAs and AAs than the other traits (especially than the lipid traits). We also obtained far smaller values of effect correlation with randomly chosen markers than with previously published GWAS. (Figure S11). These results reinforce the idea that QTL have greater similarity of effects between EAs and AAs (lower genetic heterogeneity), whereas random markers seem to have different effects in different populations (greater genetic heterogeneity).

Finally, we found differences in the estimated proportion of non-zero effects between EAs and AAs for HDL, LDL, and serum urate but not for height, reinforcing that the genetic architecture of height may be more similar between EAs and AAs in comparison to the other three traits (Figure 5 and S9). The proportion of non-zero effects markedly decreased with the -logP value threshold implying that large-effect QTL have a greater proportion of non-zero effects than small-effect QTL, especially for lipid traits. This trend is largely driven by the proportion of non-zero *main* effects for both ethnic groups (i.e. effects common to both ethnic groups – Figure S9). Finally, we also observed greater variability in posterior correlation of effects among lipid traits and serum urate in comparison to height (Figure 6).

Since we “pruned” our SNP subsets to minimize the influence of LD (see methods), other factors such as epistasis and genetic-by-environmental interaction (G×E) could explain the presence of effect heterogeneity for the traits considered in this study. Indeed, if ethnicity correlates with lifestyle, diet, income and other factors that may induce (G×E), then SNP effects can become population-specific. The estimated proportion of non-zero effects for height was in general higher than that of LDL, HDL and serum urate: three traits that are more affected by diet and lifestyle. Likewise, epistasis may be responsible for the vast majority of small additive effects (Mackay and Moore 2014) and previous studies have attributed the non-replication of genetic associations in different populations to epistasis (Greene *et al.* 2009b). Thus, epistatic gene action can also have a role in explaining differences in the allelic substitution effects of SNPs and can consequently induce effect heterogeneity (observed in our study as lower proportion of non-zero effects for small-effect QTL), especially if allele frequencies vary between the two populations (Mackay and Moore 2014; Wei *et al.* 2014) (Figure S12).

In **conclusion**, we have proposed a versatile methodology based on random-effects interactions that can apply non-Gaussian priors to marker effects for quantifying the extent of effect heterogeneity between ethnically diverse groups using a combination of variable selection and shrinkage. This proposed approach can yield estimates of SNP-heritability, average correlation of effects, proportion of non-zero effects as well as SNP-specific attributes in genomic regions of interest. Of the traits considered in our study, effect heterogeneity was lower for height than for traits influenced by lifestyle. We postulate that differences in allele frequency and in LD patterns, together with epistasis and G×E can contribute to effect heterogeneity between AAs and EAs. Further research on interactions between genes and proteins as well as signaling pathways is needed to understand the functional mechanisms underlying effect heterogeneity between ethnic groups for complex human traits and diseases.

## Acknowledgments

GDLC and YV acknowledge financial support from NIH grants GM R01099992 and GM R01101219. The authors acknowledge valuable comments provided Drs. Sadeep Shrestha, Trudy Mackay, Edward Buckler, Marguerite Irvin and Nianjun Liu. The authors also acknowledge the help of Cristen Willer and Sebanti Sengupta for providing us summary statistics for HDL and LDL from the GLGC consortium after excluding the ARIC cohort as well as Christian Gieger and Jürgen Riegel for providing summary-statistics for serum urate from the GUGC consortium after excluding the ARIC cohort.

## Literature Cited

Astle W., Balding D. J., 2009 Population Structure and Cryptic Relatedness in Genetic Association Studies. Stat. Sci. 24: 451–471.

Barnholtz-Sloan J. S., Raska P., Rebbeck T. R., Millikan R. C., 2011 Replication of GWAS “Hits” by Race for Breast and Prostate Cancers in European Americans and African Americans. Front. Genet. 2: 37.

Brant S. R., Okou D. T., Simpson C. L., Cutler D. J., Haritunians T., et al., 2017 Genome-Wide Association Study Identifies African-Specific Susceptibility Loci in African Americans With Inflammatory Bowel Disease. Gastroenterology 152: 206–217.e2.

Brown B. C., Ye C. J., Price A. L., Zaitlen N., Zaitlen N., 2016 Transethnic Genetic-Correlation Estimates from Summary Statistics. Am. J. Hum. Genet. 99: 76–88.

Bulik-Sullivan B. K., Loh P.-R., Finucane H. K., Ripke S., Yang J., et al., 2015 LD Score regression distinguishes confounding from polygenicity in genome-wide association studies. Nat. Genet. 47: 291–295.

Chen G., Doumatey A. P., Zhou J., Lei L., Bentley A. R., et al., 2017 Genome-wide analysis identifies an african-specific variant in SEMA4D associated with body mass index. Obesity (Silver Spring). 25: 794–800.

Christensen O. F., Madsen P., Nielsen B., Su G., 2014 Genomic evaluation of both purebred and crossbred performances. Genet. Sel. Evol. 46: 23.

Cockerham C., 1969 Variance of gene frequencies. Evolution (N. Y).: 72–84.

de Candia T. R., Lee S. H., Yang J., Browning B. L., Gejman P. V., et al., 2013 Additive Genetic Variation in Schizophrenia Risk Is Shared by Populations of African and European Descent. Am. J. Hum. Genet. 93: 463–470.

Deng H. W., 2001 Population admixture may appear to mask, change or reverse genetic effects of genes underlying complex traits. Genetics 159: 1319–23.

Franceschini N., Fox E., Zhang Z., Edwards T. L., Nalls M. A., et al., 2013 Genome-wide association analysis of blood-pressure traits in African-ancestry individuals reveals common associated genes in African and non-African populations. Am. J. Hum. Genet. 93: 545–54.

Gabriel S. B., 2002 The Structure of Haplotype Blocks in the Human Genome. Science (80-.). 296: 2225–2229.

Gaggiotti O. E., Bekkevold D., Jørgensen H. B. H., Foll M., Corvalho G. R., et al., 2009 Disentangling the effects of evolutionary, demographic, and environmental factors influencing genetic structure of natural populations: Atlantic herring as a case study. Evolution 63: 2939–51.

García-Cortés L., Toro M., 2006 Multibreed analysis by splitting the breeding values. Genet. Sel.

Gianola D., los Campos G. de, Hill W. G., Manfredi E., Fernando R., 2009 Additive genetic variability and the Bayesian alphabet. Genetics 183: 347–63.

Greene C. S., Penrod N. M., Williams S. M., Moore J. H., 2009a Failure to replicate a genetic association may provide important clues about genetic architecture. PLoS One 4: e5639.

Greene C. S., Penrod N. M., Williams S. M., Moore J. H., 2009b Failure to Replicate a Genetic Association May Provide Important Clues About Genetic Architecture (TIA Sorensen, Ed.). PLoS One 4: e5639.

Gudbjartsson D. F., Arnar D. O., Helgadottir A., Gretarsdottir S., Holm H., et al., 2007 Variants conferring risk of atrial fibrillation on chromosome 4q25. Nature 448: 353–7.

Habier D., Fernando R. L., Kizilkaya K., Garrick D. J., 2011 Extension of the Bayesian alphabet for genomic selection. BMC Bioinformatics 12: 186.

Haga S. B., 2010 Impact of limited population diversity of genome-wide association studies. Genet. Med. 12: 81–4.

Ishwaran H., Rao J. S., 2005 Spike and slab variable selection: Frequentist and Bayesian strategies. Ann. Stat. 33: 730–773.

Karoui S., Corabaño M. J., Díaz C., Legarra A., VanRaden P., et al., 2012 Joint genomic evaluation of French dairy cattle breeds using multiple-trait models. Genet. Sel. Evol. 44: 39.

Kraft P., Zeggini E., Ioannidis J. P. A., 2009 Replication in genome-wide association studies. Stat. Sci. 24: 561–573.

Lander E. S., Schork N. J., 1994 Genetic dissection of complex traits. Science 265: 2037–48.

Lee S. H., Yang J., Goddard M. E., Visscher P. M., Wray N. R., 2012 Estimation of pleiotropy between complex diseases using single-nucleotide polymorphism-derived genomic relationships and restricted maximum likelihood. Bioinformatics 28: 2540–2.

Lehermeier C., Schön C.-C., los Campos G. de, 2015 Assessment of Genetic Heterogeneity in Structured Plant Populations Using Multivariate Whole-Genome Regression Models. Genetics 201.

Lehermeier C., los Campos G. de, Wimmer V., Schön C.-C., 2017 Genomic variance estimates: With or without disequilibrium covariances? J. Anim. Breed. Genet. 134: 232–241.

Li Y. R., Keating B. J., 2014 Trans-ethnic genome-wide association studies: advantages and challenges of mapping in diverse populations. Genome Med. 6: 91.

Liu N., Zhao H., Patki A., Limdi N. A., Allison D. B., 2011 Controlling Population Structure in Human Genetic Association Studies with Samples of Unrelated Individuals. Stat. Interface 4: 317–326.

los Campos G. de, Naya H., Gianola D., Crossa J., Legarra A., et al., 2009 Predicting quantitative traits with regression models for dense molecular markers and pedigree. Genetics 182: 375–85.

los Campos G. de, Hickey J., Pong-Wong R., Daetwyler H., 2013 Whole-genome regression and prediction methods applied to plant and animal breeding. Genetics 193: 327–345.

los Campos G. de, Sorensen D., Gianola D., 2014 Genomic Heritability: What Is It? 10th World Congr. Genet. Appl. to Livest. Prod.

los Campos G. de, Sorensen D., 2014 On the genomic analysis of data from structured populations. J. Anim. Breed. Genet. 131: 163–4.

los Campos G. de, Veturi Y., Vazquez A. I., Lehermeier C., Pérez-Rodríguez P., 2015 Incorporating Genetic Heterogeneity in Whole-Genome Regressions Using Interactions. J. Agric. Biol. Environ. Stat. 20: 467–490.

Mackay T. F., Moore J. H., 2014 Why epistasis is important for tackling complex human disease genetics. Genome Med. 6: 124.

Malécot G., 1947 Les Mathématiques de l’hérédité. Masson, Paris.

Marchini J., Cordon L. R., Phillips M. S., Donnelly P., 2004 The effects of human population structure on large genetic association studies. Nat. Genet. 36: 512–7.

Marigorta U. M., Navarro A., 2013 High Trans-ethnic Replicability of GWAS Results Implies Common Causal Variants (SM Williams, Ed.). PLoS Genet. 9: e1003566.

Meuwissen T. H., Hayes B. J., Goddard M. E., 2001 Prediction of total genetic value using genome-wide dense marker maps. Genetics 157: 1819–29.

Ng M. C. Y., Shriner D., Chen B. H., Li J., Chen W.-M., et al., 2014 Meta-Analysis of Genome-Wide Association Studies in African Americans Provides Insights into the Genetic Architecture of Type 2 Diabetes (E Zeggini, Ed.). PLoS Genet. 10: e1004517.

Ntzani E. E., Liberopoulos G., Manolio T. A., Ioannidis J. P. A., 2012 Consistency of genome-wide associations across major ancestral groups. Hum. Genet. 131: 1057–1071.

Okada Y., Wu D., Trynka G., Raj T., Terao C., et al., 2014 Genetics of rheumatoid arthritis contributes to biology and drug discovery. Nature 506: 376–81.

Olson K. M., VanRaden P. M., Tooker M. E., 2012 Multibreed genomic evaluations using purebred Holsteins, Jerseys, and Brown Swiss. J. Dairy Sci. 95: 5378–5383.

Omori S., Tanaka Y., Takahashi A., Hirose H., Kashiwagi A., et al., 2008 Association of CDKAL1, IGF2BP2, CDKN2A/B, HHEX, SLC30A8, and KCNJ11 with susceptibility to type 2 diabetes in a Japanese population. Diabetes 57: 791–5.

Park T., Casella G., 2008 The Bayesian Lasso. J. Am. Stat. Assoc. 103: 681–686.

Park S. L., Cheng I., Haiman C. A., 2017 Genome-wide association studies of cancer in diverse populations.

Peprah E., Xu H., Tekola-Ayele F., Royal C. D., 2015 Genome-wide association studies in Africans and African Americans: expanding the framework of the genomics of human traits and disease. Public Health Genomics 18: 40–51.

Pérez P., los Campos G. de, 2014 Genome-wide regression and prediction with the BGLR statistical package. Genetics 198: 483–95.

Pfenninger M., Salinger M., Haun T., Feldmeyer B., 2011 Factors and processes shaping the population structure and distribution of genetic variation across the species range of the freshwater snail radix balthica (Pulmonata, Basommatophora). BMC Evol. Biol. 11: 135.

Prasad S., Bhatia T., Kukshal P., Nimgaonkar V. L., Deshpande S. N., et al., 2017 Attempts to replicate genetic associations with schizophrenia in a cohort from north India. npj Schizophr. 3: 28.

Price A. L., Zaitlen N. A., Reich D., Patterson N., 2010 New approaches to population stratification in genome-wide association studies. Nat. Rev. Genet. 11: 459–463.

Puckett E. E., Kristensen T. V, Wilton C. M., Lyda S. B., Noyce K. V, et al., 2014 Influence of drift and admixture on population structure of American black bears (Ursus americanus) in the Central Interior Highlands, USA, 50 years after translocation. Mol. Ecol. 23: 2414–27.

Reeve E., 1953 Studies in quantitative inheritance. 3. Heritability and genetic correlation in progeny tests using different mating systems - Google Scholar. J. Genet. 51: 520–542.

Rosenberg N. A., Huang L., Jewett E. M., Szpiech Z. A., Jankovic I., et al., 2010 Genome-wide association studies in diverse populations. Nat. Rev. Genet. 11: 356–66.

Shifman S., 2003 Linkage disequilibrium patterns of the human genome across populations. Hum. Mol. Genet. 12: 771–776.

Speed D., Balding D. J., Dk D. A., 2018 Exposing flaws in S-LDSC; reply to Gazal et al.

Taylor J. Y., Schwander K., Kardia S. L. R., Arnett D., Liang J., et al., 2016 A Genome-wide study of blood pressure in African Americans accounting for gene-smoking interaction. Sci. Rep. 6: 18812.

Tsai E. A., Grochowski C. M., Loomes K. M., Bessho K., Hakonarson H., et al., 2014 Replication of a GWAS signal in a Caucasian population implicates ADD3 in susceptibility to biliary atresia. Hum. Genet. 133: 235–43.

Vlaming R. de, Okbay A., Rietveld C. A., Johannesson M., Magnusson P. K. E., et al., 2017 Meta-GWAS Accuracy and Power (MetaGAP) Calculator Shows that Hiding Heritability Is Partially Due to Imperfect Genetic Correlations across Studies (J Marchini, Ed.). PLOS Genet. 13: e1006495.

Wei M., Werf J. Van der, 1994 Maximizing genetic response in crossbreds using both purebred and crossbred information. Anim. Prod.

Wei W.-H., Hemani G., Haley C. S., 2014 Detecting epistasis in human complex traits. Nat. Rev. Genet. 15: 722–33.

Welter D., MacArthur J., Morales J., Burdett T., Hall P., et al., 2014 The NHGRI GWAS Catalog, a curated resource of SNP-trait associations. Nucleic Acids Res. 42: D1001–6.

Wright S., 1949 The Genetical Structure of Populations. Ann. Eugen. 15: 323–354.

Yamada H., Penney K. L., Takahashi H., Katoh T., Yamano Y., et al., 2009 Replication of prostate cancer risk loci in a Japanese case-control association study. J. Natl. Cancer Inst. 101: 1330–6.

Zhou X., Cheung C.-L., Karasugi T., Karppinen J., Samartzis D., et al., 2018 Trans-ethnic polygenic analysis supports genetic overlaps of lumbar disc degeneration with height, body mass index, and bone mineral density. biorxiv.

